# MobsPy: A Meta-Species Language for Chemical Reaction Networks*

**DOI:** 10.1101/2022.05.05.490768

**Authors:** Fabricio Cravo, Matthias Függer, Thomas Nowak, Gayathri Prakash

## Abstract

Chemical reaction networks are widely used to model biochemical systems. However, when the complexity of these systems increases, the chemical reaction networks are prone to errors in the initial modeling and subsequent updates of the model.

We present the Meta-species-oriented Biochemical Systems Language (MobsPy), a language designed to simplify the definition of chemical reaction networks in Python. MobsPy is built around the notion of meta-species, which are sets of species that can be multiplied to create higher-dimensional orthogonal characteristics spaces and inheritance of reactions. Reactions can modify these characteristics. For reactants, queries allow to select a subset from a meta-species and use them in a reaction. For products, queries specify the dimensions in which a modification occurs. We demonstrate the simplification capabilities of the MobsPy language at the hand of a running example and a circuit from literature. The MobsPy Python package includes functions to perform both deterministic and stochastic simulations, as well as easily configurable plotting. The MobsPy package is indexed in the Python Package Index and can thus be installed via `pip`.

## 1 Introduction

Chemical reaction networks (CRNs) model the dynamics of biochemical systems by a set of species and reactions acting on them [8]. In particular, in synthetic biology, which studies the engineering of new behavior in biological entities [10], CRNs have proved useful to model genetic circuits. Examples are the toggle switch, the repressilator, and logic gates, among others [16,3,1]. While synthetic biology has primarily addressed single-cell behavior, recent work has studied the engineering of bacterial populations to reduce cell burden and allow for population-averaging of circuit responses [15,17].

However, the modeling of genetic circuits using CRNs is error-prone due to the complexity of the resulting model. Lopez, Muhlich, Bachman, and Sorger [13] studied several works from literature and found discrepancies when comparing their description and the provided models. There is thus a need for tools that simplify the modeling of complex biological circuits through CRNs, both for single-cell and population-level dynamics.

Several solutions have been proposed to facilitate the modeling and simulation of such systems. For instance, COPASI [11] and iBioSim [14] are simulators that can be either used via a graphical interface or by writing a model in SBML format [12]. They come with state-of-the-art stochastic and ordinary differential equation (ODE) solvers, but are not directly suited for automated prototyping due to design entry by a GUI. BasiCO [2] is a Python interface for using COPASI, thus allowing for automation. However, BasiCO does not support any language-based model simplifications: species and reactions are added one by one. BSim [9] is a geometric, agent-based simulator for population dynamics and uses ODEs for internal cellular dynamics.

In this work, we propose MobsPy, a language and Python framework that is based on the concept of meta-species and meta-reactions. Meta-species are sets of species and meta-reactions act upon meta-species. This allows for model simplification via the following features: (i) *Inheritance of reactions.* Meta-reactions construct reactions for all the species that inherit from the meta-species present in the meta-reaction. (ii) *The definition of species via independent characteristics-space structures.* Products of meta-species create a meta-species whose char-acteristics-space is the Cartesian product of the characteristics-space of its elements. (iii) *The possibility to query and transform the characteristics of a species.* Queries can be used to restrict meta-species to subsets of species. Transformations can be used to change specific characteristics within a meta-species’ characteristics-space. These features will be discussed in greater detail in Section 2, along with an example model.

The most closely related frameworks to Mobspy are Kappa [6] and iBioGen [7], both providing a rule-based language that can define multiple reactions with a single rule. A rule defines how individual species (typically molecules) combine and separate based on their state. While Kappa was designed to focus on complex formation in chemical pathways, MobsPy targets more general dynamics, where the reacting species do not necessarily form complexes upon interaction but may change states, produce, or consume species. Kappa also proposes the utilization of gadgets for keeping track of complex state changes, while MobsPy uses Python itself. In [5], an extension with general inheritance has been proposed to Kappa. To achieve this, they propose a distinction between two types of agents called concrete and generic agents, which is not required for inheritance in MobsPy. Also, Kappa’s inheritance extension does not allow for multiple inheritances.

## 2 MobsPy syntax and simulator

We next discuss the main features of MobsPy along with a running example. Consider a system of two groups of trees (Figure 1a): one group where trees grow and reproduce in a dense population and one in a sparse population. We assume that the leaves of a tree change color from green to yellow to brown in a cyclic fashion. Further, trees die, with young trees in the dense group dying at a higher rate due to competition for resources and space.

**Fig. 1:**
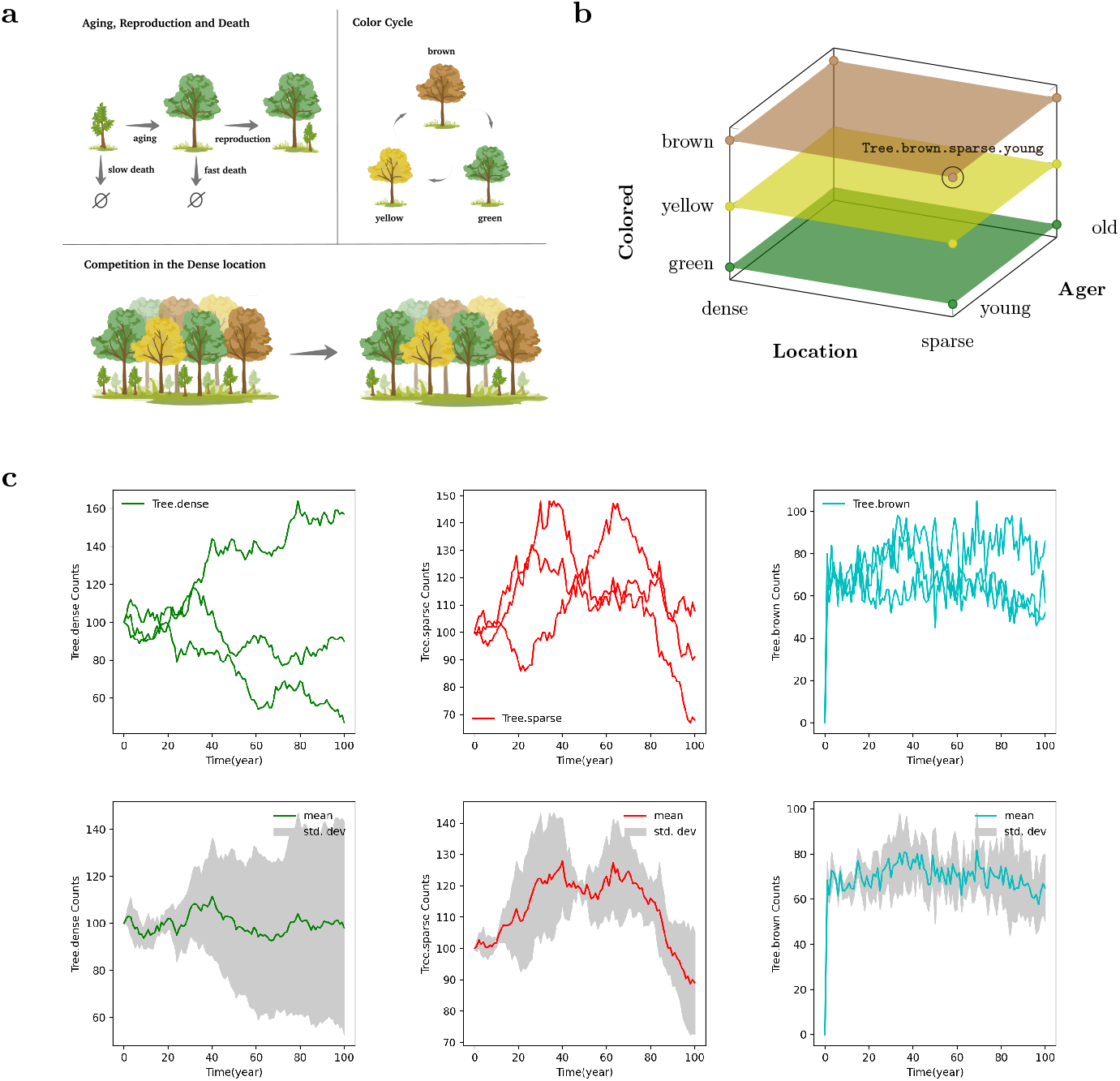
**(a)** System dynamics: schematic representation of all the reactions in the Tree model, which are aging, reproduction, color cycle, and competition. **(b)** Meta-species Tree created by multiplication of the three base-species Ager, Location, and Colored. One of the twelve species generated, Tree.brown.sparse.young, has been labeled. **(c)** MobsPy default plots after simulating the Tree model (n = 3 stochastic runs). In the top row of the panel individual runs are shown. The bottom plots depicts mean and standard deviation.

We start modeling the system by defining base-species with characteristics, and a reaction for how the base-species Ager changes its characteristics:

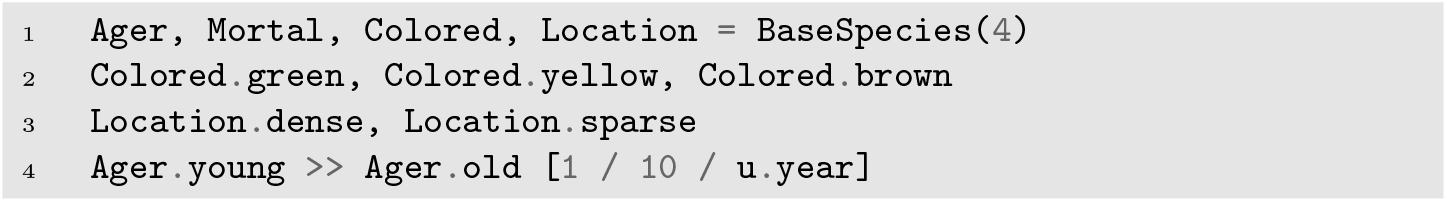

A base species has a single set of characteristics that can either be defined explicitly using the dot notation as in code line 2 or implicitly as in code line 4. In the implicit case, new characteristics are automatically added, when they are used for the first time inside a reaction.

*Meta-reactions* have a CRN-like syntax of the form `reactants >> products [rate_spec]`. The `rate_spec` can be a (non-negative) real or a function whose parameters are the meta-species reactants, and that returns a real or a string. In the case of a (returned) real, the reaction rate follows mass-action kinetics, with the real being the rate constant. In the case of a returned string, one can define different kinetics in terms of concentrations of species/characteristics of the reactants.

To assign different death rates for old and young trees, we make use of a `rate_spec` in terms of a function that returns the respective rate-constant (see code line 5), and the special base-species `Zero` that contains no species.

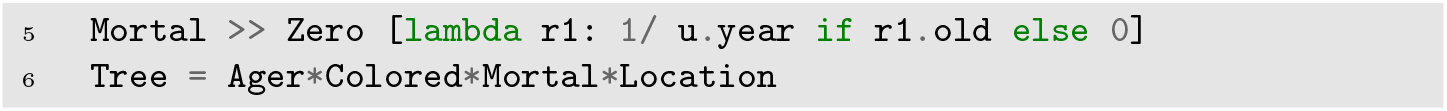

*Multiplication* (see code line 6) is used to combine base-species and meta-species into more complex meta-species. The product meta-species inherits all characteristics from its factors. In our example, `Tree` contains twelve species (see Figure 1b). Further, the product meta-species inherits the meta-reactions of its factors. Therefore, `Tree` receives the aging reactions from `Ager` and the death reactions from `Mortal`. This allows to significantly simplify the model: only one death and aging meta-reaction need to be specified for all the model’s species.

Within reactants, the *dot operator* is used to query for species inside meta-species, making it possible to assign reactions to only the preferred sub-set. Queries can be composed arbitrarily and independently of order. It is further possible to query over a string value stored inside a variable, say s, using `species.c(s)`. In our example, we make use of queries to specify the color-cycle exhibited by the leaves:

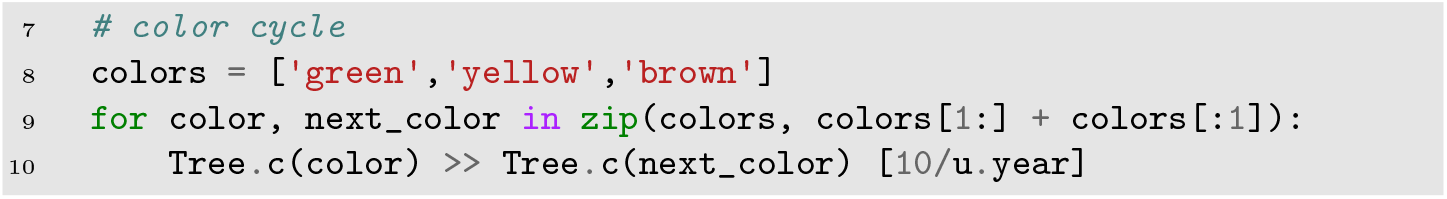

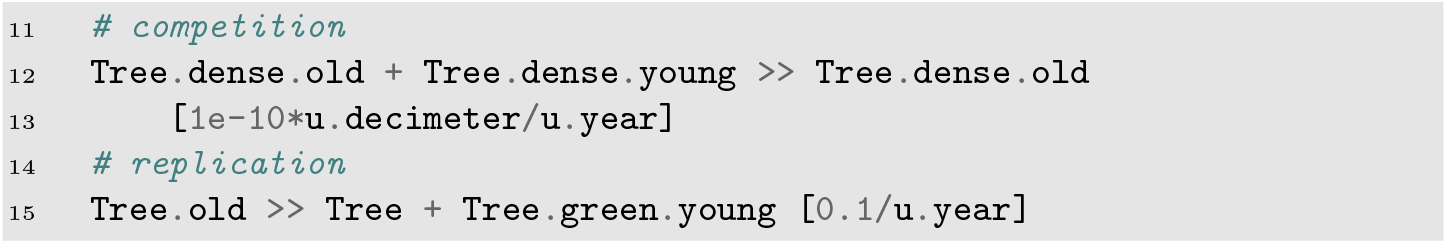

Unlike in reactions of a CRN, *order of products and reactants* matters in meta-reactions: In the competition meta-reaction (code line 12), `Tree.old` wins the competition against `Tree.young` and becomes `Tree.old`. MobsPy, by default, uses a round-robin order, where products cycle through the list of available reactants from the same meta-species. Alternatively, labels can be used to explicitly declare which reactant becomes which product.

The *initial count* of the default species within a meta-species `S` is set via `S`(*i*), where *i* is either an integer count, or a real-valued concentration with respective units. Here, the default species is the species with the first characteristic added to each meta-species from which it inherits. For example, in the Tree model, `Tree(50)` assigns the count 50 to `Tree.green.dense.young`. To assign counts to other species, one can add more characteristics.

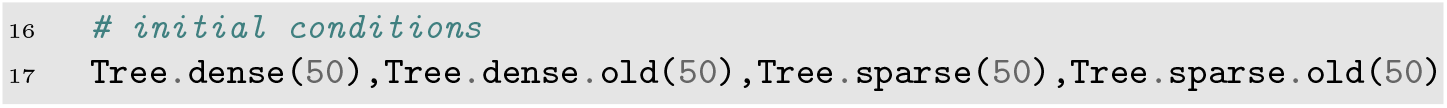

A simulation object of the species in the meta-species `Tree` is finally constructed by:

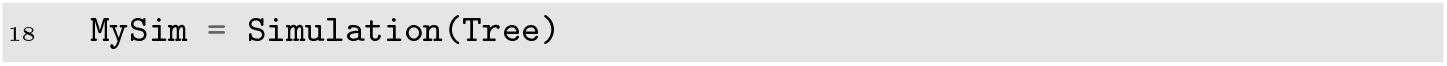

The simulation can then be parametrized with solver parameters (stochastic and ODE), plotting options, and data export options. As a backend, Mobspy generates SBML models [12] and runs simulations using BasiCO. Simulation results can be accessed directly in Python or exported as JSON files. Examples for default stochastic plots are shown in Figure 1c. The plots can be easily configured via a Python dictionary, where a single parameter can be set for all figures, for individual figures, or for each curve in a hierarchical fashion. For a detailed description of the modeling, simulation, and plotting features of MobsPy, we refer the reader to the GitHub repository [4]. It contains several examples like a simple reaction `A + B >> 2*C + D`, a Hill function model of a genetic oscillator, a bacteria-phage system, a bacteria and phage random walk, a simple repressor, a toggle switch, genetic gates such as the `NOR` and `AND` gates, an example for the hierarchical plot structure and tutorial models.

In the next section, we demonstrate the modeling capabilities of MobsPy at the hand of a real-world synthetic design from literature [3].

## 3 Genetic circuits with MobsPy: the CRISPRlator

To show that MobsPy is well-suited to model genetic circuits, we modeled the CRISPRlator, a CRISPRi oscillator proposed by Santos-Moreno, Tasiudi, Stelling, and Schaerli [16]. The circuit is a CRISPRi-based repressilator where each gRNA represses the next gRNA-encoding gene in a cyclic manner. The model is shown below, with reaction rates taken from Clamons and Murray [3].

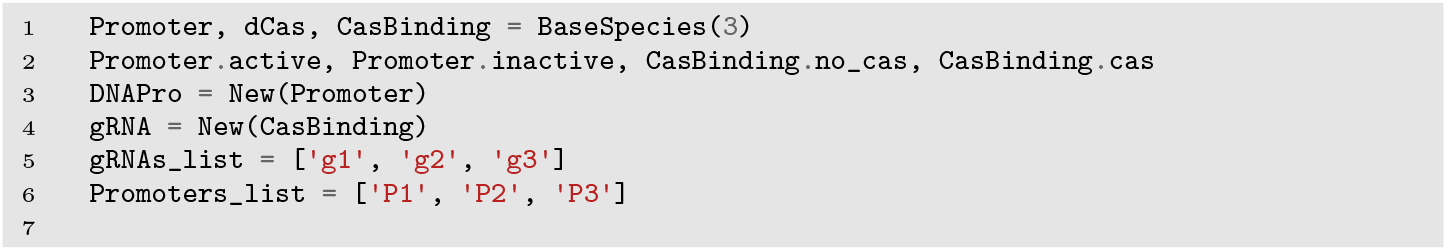
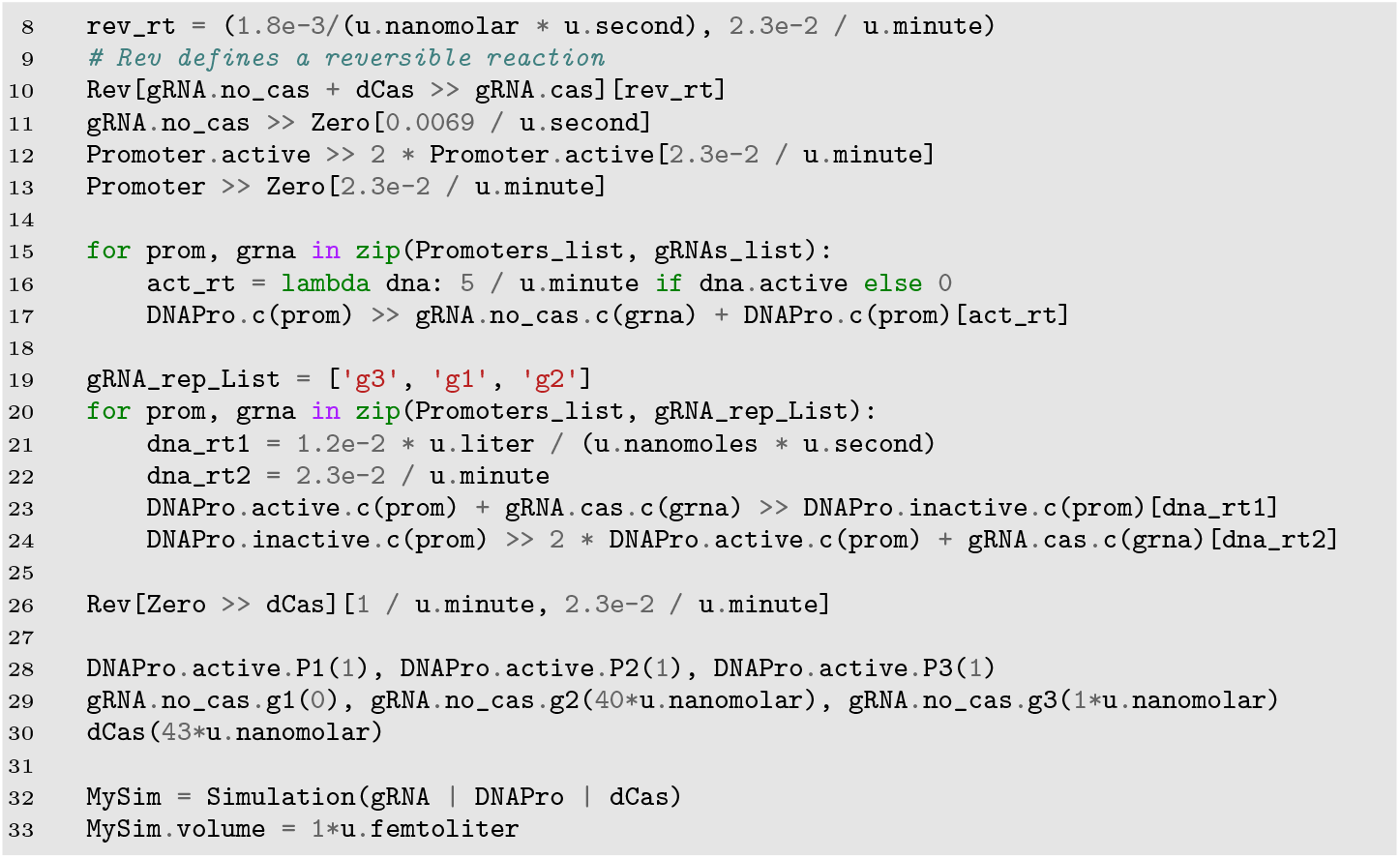

The code to run ODE simulations over a range of 650 h is shown in Figure 2a. The resulting plot is depicted in Figure 2b.

**Fig. 2:**
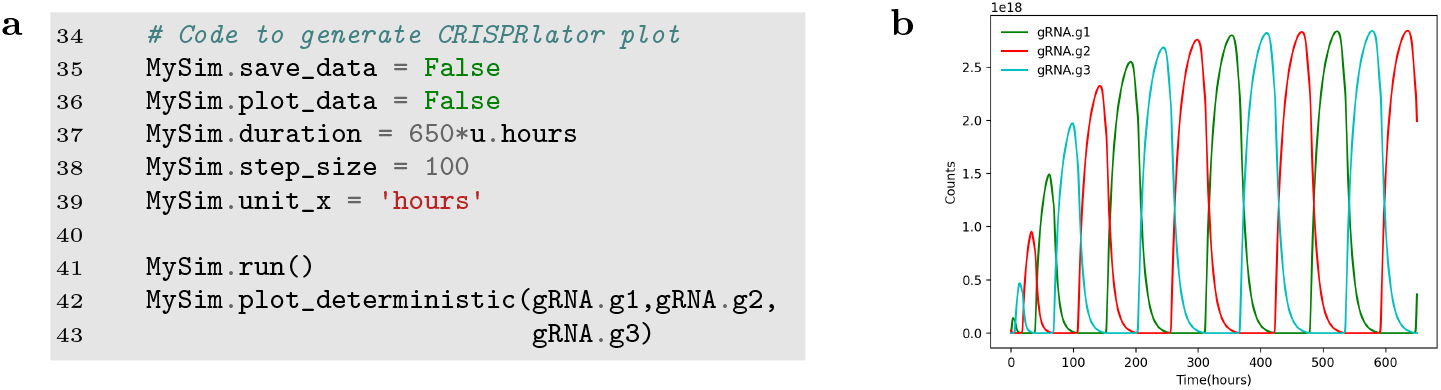
**(a)** Code for simulating the CRISPRlator model and generating default plots. **(b)** CRISPRlator simulation results.

## 4 Conclusions

We presented MobsPy, a simple CRN programming language to simplify the modeling of complex biochemical reaction networks. The simulator creates SBML code and uses BasiCO/COPASI as a stochastic and ODE solver. We discussed MobsPy’s features at the hand of a Tree toy example and a real-world genetic design, with succinct model descriptions instead of a direct specification as CRNs (12 species and 45 reactions for the Tree and 13 species and 32 reactions for the CRISPRi model). MobsPy is open-source (MIT license) and easy to install through pip or via the git repository.

## References

1. Adam Arkin and John Ross. Computational functions in biochemical reaction networks. Biophysical Journal, 67(2):560–578, 1994.

2. Frank T. Bergmann. BasiCO. https://github.com/copasi/basico, 2022.

3. Samuel E. Clamons and Richard M. Murray. Modeling dynamic transcriptional circuits with CRISPRi. https://www.biorxiv.org/content/early/2017/11/27/22531, 2017.

4. Fabricio Cravo, Matthias Függer, Thomas Nowak, and Gayathri Prakash. MobsPy. https://github.com/ROBACON/mobspy, 2022.

5. Vincent Danos, Jérôme Feret, Walter Fontana, Russ Harmer, and Jean Krivine. Rule-based modelling and model perturbation. In Transactions on Computational Systems Biology XI, volume 5750 of Lecture Notes in Bioinformatics, pages 116–137. Springer, Heidelberg, 2009.

6. Vincent Danos and Cosimo Laneve. Formal molecular biology. Theoretical Computer Science, 325(1):69–110, 2004.

7. James R. Faeder, Michael L. Blinov, and William S. Hlavacek. Rule-based modeling of biochemical systems with BioNetGen. In Ivan V. Maly, editor, Systems Biology. Methods in Molecular Biology (Methods and Protocols), volume 500 of Methods in Molecular Biology, pages 113–167. Springer Science+Business Media, New York, 2009.

8. Daniel T. Gillespie. Exact stochastic simulation of coupled chemical reactions. The Journal of Physical Chemistry, 81(25):2340–2361, 1977.

9. Thomas E. Gorochowski, Antoni Matyjaszkiewicz, Thomas Todd, Neeraj Oak, Kira Kowalska, Stephen Reid, Krasimira T. Tsaneva-Atanasova, Nigel J. Savery, Claire S. Grierson, and Mario Di Bernardo. BSim: an agent-based tool for modeling bacterial populations in systems and synthetic biology. PLoS ONE, 7(8):e42790, 2012.

10. Matthias Heinemann and Sven Panke. Synthetic biology—putting engineering into biology. Bioinformatics, 22(22):2790–2799, 2006.

11. Stefan Hoops, Sven Sahle, Ralph Gauges, Christine Lee, Jürgen Pahle, Natalia Simus, Mudita Singhal, Liang Xu, Pedro Mendes, and Ursula Kummer. COPASI-a COmplex PAthway SImulator. Bioinformatics, 22(24):3067–3074, 2006.

12. M. Hucka, A. Finney, H. M. Sauro, H. Bolouri, J. C. Doyle, H. Kitano, A. P. Arkin, B. J. Bornstein, D. Bray, A. Cornish-Bowden, A. A. Cuellar, S. Dronov, E. D. Gilles, M. Ginkel, V. Gor, I. I. Goryanin, W. J. Hedley, T. C. Hodgman, J.-H. Hofmeyr, P. J. Hunter, N. S. Juty, J. L. Kasberger, A. Kremling, U. Kummer, N. Le Novère, L. M. Loew, D. Lucio, P. Mendes, E. Minch, E. D. Mjolsness, Y. Nakayama, M. R. Nelson, P. F. Nielsen, T. Sakurada, J. C. Schaff, B. E. Shapiro, T. S. Shimizu, H. D. Spence, J. Stelling, K. Takahashi, M. Tomita, J. Wagner, J. Wang, and the rest of the SBML Forum. The systems biology markup language (SBML): a medium for representation and exchange of biochemical network models. Bioinformatics, 19(4):524–531, 2003.

13. Carlos F. Lopez, Jeremy L. Muhlich, John A. Bachman, and Peter K. Sorger. Programming biological models in Python using PySB. Molecular Systems Biology, 9:646, 2013.

14. Chris J. Myers, Nathan Barker, Kevin Jones, Hiroyuki Kuwahara, Curtis Madsen, and Nam-Phuong D. Nguyen. iBioSim: a tool for the analysis and design of genetic circuits. Bioinformatics, 25(21):2848–2849, 2009.

15. Sergi Regot, Javier Macia, Núria Conde, Kentaro Furukawa, Jimmy Kjellén, Tom Peeters, Stefan Hohmann, Eulãlia De Nadal, Francesc Posas, and Ricard Solé. Distributed biological computation with multicellular engineered networks. Nature, 469(7329):207–211, 2011.

16. Javier Santos-Moreno, Eve Tasiudi, Joerg Stelling, and Yolanda Schaerli. Multistable and dynamic CRISPRi-based synthetic circuits. Nature Communications, 11(1):1–8, 2020.

17. Alvin Tamsir, Jeffrey J. Tabor, and Christopher A. Voigt. Robust multicellular computing using genetically encoded NOR gates and chemical ‘wires’. Nature, 469(7329):212–215, 2011.

